# A confined gene drive for population modification in the malaria vector *Anopheles stephensi*

**DOI:** 10.64898/2026.04.01.715791

**Authors:** Xuejiao Xu, Yiran Liu, Xihua Jia, Jie Yang, Yunchen Xia, Jingheng Chen, Jackson Champer

## Abstract

Gene drives are genetic elements that bias their own inheritance to spread desired traits in target populations, enabling population modification or suppression. Although homing-based drives can propagate efficiently, their potential for uncontrolled spread may present a challenge for field deployment. Thus, confined drive systems are needed. Here, we developed a confined modification drive, called Toxin-Antidote Recessive Embryo (TARE) drive, in the globally important malaria vector *Anopheles stephensi*. This drive works by cleaving and disrupting wild-type alleles in the germline or early embryo from maternally deposited Cas9. Disrupted alleles are recessive lethal, thus increasing the drive in a frequency-dependent manner. Inheritance bias was moderate in crosses between drive heterozygote mosquitoes, possibly due to low gRNA activity and thus moderate germline cleavage rates. Single-release cage trials confirmed the TARE drive’s ability to spread, although the drive ultimately declined due to fitness costs and resistance alleles associated with repetitive elements. Nonetheless our modelling analysis indicate that this TARE system could achieve population spread if the resistance issue is addressed. These findings demonstrate a functional prototype TARE drive in *Anopheles stephensi* and highlight key parameters governing its performance. Minor design optimizations could substantially improve efficiency and integrity, enabling rapid but confined population modification.

## Introduction

“Selfish genes” are naturally existing alleles that can bias their own inheritance, allowing them to spread through a population at rates exceeding the Mendelian expectation. Building on this concept, synthetic gene drive systems have been proposed, introducing these elements into target organisms(Nolan 2021, Bier 2022, Wang et al. 2025). Such engineered systems can be designed for population suppression by disrupting genes essential for reproduction or viability or by biasing the sex ratio. They can also be designed for population modification by introducing beneficial traits such as pathogen transmission reduction(Carballar-Lejarazú et al. 2023) or pesticide sensitivity(Guichard et al. 2019) into target populations.

Compared with traditional chemical-based pest management strategies, gene drives offer advantages including persisting in populations across generations, acting with high species specificity, and avoiding environmental impacts associated with insecticide use, thereby attracting increasing attention from researchers and policy makers. Despite their promise, development of engineered gene drive systems was slow until the emergence of CRISPR/Cas9 editing technology, which provides an efficient and programmable means of manipulating target genomic sequences. This technical breakthrough greatly expanded the feasibility and flexibility of gene drive designs across various organisms. Since then, they have been demonstrated in several species including yeast(DiCarlo et al. 2015), mice(Grunwald et al. 2019, Pfitzner et al. 2020), viruses(Walter et al. 2024), plants(Liu et al. 2024, Oberhofer et al. 2024), and insect species(Champer et al. 2020d, Faber et al. 2024, Kyrou et al. 2018, Carballar-Lejarazú et al. 2023, Xu et al. 2025, Yadav et al. 2023, Xu et al. 2022, Pescod et al. 2024, Anderson et al. 2023, Speth et al. 2025, Yang et al. 2025, Larrosa-Godall et al. 2025).

The most widely studied gene drives rely on a homing mechanism with CRISPR/Cas9(Verkuijl et al. 2022, Ang et al. 2025). In this system, the drive allele, which encodes both Cas9 and gRNA elements, expresses a Cas9/gRNA complex that cleaves the homologous wild-type allele, generating a double-strand break. This break can then be repaired by either the homology-directed repair or end-joining pathways. Homology-directed repair uses the drive allele as a repair template, copying the drive into the target locus. In contrast, end-joining often produces mutations, which can preserve the function of the target gene but usually disrupt it. Such alleles, no longer recognized by the Cas9/gRNA complex, become drive-resistant alleles. Consequently, the spread efficiency of a homing gene drive depends on the homology-directed repair frequency after drive cleavage.

A series of studies aimed to build modification drives in *Anopheles* mosquitoes by targeting eye pigmentation genes, showcasing high drive efficiency in cage populations(Gantz et al. 2015, Adolfi et al. 2020, Carballar-Lejarazú et al. 2023). However, resistance formation remained a potentially unresolved challenge, even with improved promoters or rescue elements. If such resistance alleles have a fitness advantage over the drive, they may be able to outcompete it, reducing the time that the drive could protect against pathogen transmission. However, by combining multiplexed gRNAs and a rescue element in a drive targeting an essential gene, resistance could be avoided in fruit flies, or at least kept to a manageable level(Champer et al. 2020d).

Yet, these homing drives still are copied by homology-directed repair, an error prone pathway that could lead to mutational inactivation of cargo genes. Further, homing drives are not confined to a target population, but will instead spread with even low levels of migration. To address this, the Toxin-Antidote Recessive Embryo (TARE) drive was developed in *Drosophila melanogaster*(Champer et al. 2020c). In TARE drive, Cas9/gRNA cleavage disrupts the target gene to generate lethality (the toxin), while the drive allele simultaneously provides a recoded version of target gene (the antidote). This recoded copy cannot be cleaved by the Cas9/gRNA complex and restores the function of target gene. As a result, only individuals inheriting at least one functional copy of the target gene (either a wild-type allele or the drive allele) can survive, whereas progeny inheriting two disrupted alleles are nonviable. Over successive generations, this selective pressure eliminates wild-type alleles from the population, allowing the rescue-bearing gene drive allele to increase in frequency. The same concept was also reported as Cleave and Rescue (ClvR), also with demonstrations in *D. melanogaster*(Oberhofer et al. 2019, 2020).

Compared to homing drives, TARE drives could be more practical from an engineering standpoint because they do not rely on precise homology-directed repair with the drive. This means that DNA cleavage often needs to be restricted to meiosis and thus relies on Cas9 germline expression, which can be challenging to achieve. Notably, maternal deposition of Cas9, which typically generates embryo resistance alleles and impedes the spread of homing drives, instead enhances inheritance bias in TARE systems. This relaxed requirement for precise Cas9 timing broadens the design flexibility of TARE drives. Though not an issue for *Anopheles* with high drive conversion rates, such a feature may be useful in other species, and indeed, such drives have already seen success in *Arabidopsis*(Liu et al. 2024, Oberhofer et al. 2024).

While homing drives are unconfined, the TARE drive has threshold-dependent invasion dynamics, meaning that it can only spread in a population when introduced above a certain frequency(Champer et al. 2020c, 2020b). This property provides an additional layer of confinement and makes the system more suitable for controlled or geographically limited applications. The confinement level of TARE drive may be further increased by adding another TARE system at a distant genomic location (2-locus TARE), with each drive targeting the gene that the other provides rescue for(Champer et al. 2020a). Though the main application of TARE drive is for population modification, confined population suppression could also be achieved by combining TARE with drives targeting reproduction or viability in a tethered drive configuration(Dhole et al. 2019, Metzloff et al. 2022, Feng and Champer 2024).

In this study, we constructed a confined TARE gene drive in the globally important invasive urban malaria mosquito vector *Anopheles stephensi*. The TARE construct was functional in biasing drive inheritance as well as cleaving and rescuing the target gene. However, its cut rates were limited, possibly caused by low efficiency of gRNA expression. It was able to spread in the cage, but was impeded by fitness costs and functional resistance allele formation due to repetitive DNA between the rescue element and native gene. Nevertheless, it represents a proof-of-principle for TARE drive in mosquitoes while pointing the way toward improvements for higher effectiveness.

## Methods

### Mosquito strain

The *Anopheles stephensi* TARE strain used in this study was generated previously(Xu et al. 2025) and designed similarly to a prototype system in *Drosophila melanogaster* (Champer et al. 2020c). The construct comprised a recoded sequence of the *hairy* target gene, a Cas9 coding sequence driven by the *vasa* promoter, a 3×P3-EGFP-SV40 fluorescence marker, and a multiplexed ribozyme-gRNA cassette controlled by the *A. stephensi* U6a promoter. The ribozyme-gRNA cassette contained four gRNAs (gRNA1: 5’-tccagctttgagtgacgggc-3’; gRNA2: 5’-gatgcttcaccgtcatctcc-3’; gRNA3: 5’-ttgaacttgttcatcacgct-3’; gRNA4: 5’-ccgggaagcggcccacctcc-3’) each separated by self-cleaving ribozymes. These gRNAs targeted sites located ∼600 bp downstream of the TARE insertion site to prevent homing events. PAM sites in the right homology arm of the donor construct were mutated to avoid unintended cleavage of the donor plasmid.

### Construction of *D. melanogaster* strain

The drive construct, containing a ribozyme-gRNA cassette and a 3×P3-DsRed-SV40 marker, was assembled using HiFi DNA Assembly (New England Biolabs). The ribozyme-gRNA cassette was adapted from a previously described construct targeting *yellow-G*, in which four gRNAs were linked by tRNA sequences(Yang et al. 2022). The same gRNA sequences were used here to enable direct comparison between this ribozyme-gRNA construct and the previous tRNA-gRNA one. Red fluorescence was used for screening to differentiate this line from an existing *nanos*-Cas9 line marked with green fluorescence(Du et al. 2024). Plasmids constructed for this study is available on GitHub (https://github.com/jchamper/Anopheles-TARE-Drive).

The fly embryo injection was conducted by UniHuaii Company in a *w^1118^* strain using a mixture of 500 ng/µL Cas9-expressing helper TTChsp70c9, 100 ng/µL gRNA helper TTTygU4 (targeting *yellow-G* for knock-in) and 500 ng/µL ribozyme-gRNA donor plasmids. The injected flies were maintained and crossed to *w^1118^* flies to obtain offspring that could be screened for the construct. Because homozygous females were sterile, strains were maintained as mixed populations containing homozygote, heterozygous, and wild-type individuals.

### Insect rearing

The rearing procedures for mosquitoes and flies followed those published previously(Faber et al. 2024, Xu et al. 2025). *A. stephensi* mosquitoes were maintained at 27 ± 1 °C, 75% relative humidity, under a 12-h light/dark cycle. Larvae and adults were fed with fish food (Hikari) and 10% sucrose, respectively. Adults were housed either in 30 × 30 cm cages or paper cups for pool crosses or single-female fertility assays. Females were blood-fed using a Hemotek membrane feeder with defibrinated cow blood. *D. melanogaster* stocks were maintained in a 25 °C incubator with a 14/10-h day/night cycle, and reared on Cornell standard cornmeal medium modified by increasing agar amount to 10 g, adding 5 g soy flour, and removing the phosphoric acid.

### Crosses and phenotypes

For *A. stephensi*, drive efficiency was assessed by crossing heterozygous transgenic mosquitoes either with siblings or with wild-type individuals. Progeny were screened at the larval stage for EGFP fluorescence, and the number of drive carriers and non-carriers were recorded.

For *D. melanogaster*, the ribozyme-gRNA line was crossed with a Cas9 line (SNc9NG) to generate double heterozygotes. These were then crossed to *w^1118^* to assess drive inheritance. Adult progeny were subsequently screened for red fluorescence (drive allele) and green fluorescence (Cas9 allele). For phenotyping, flies were anaesthetized with CO₂ and screened using the NIGHTSEA system.

To assess fertility of TARE heterozygotes, five homozygous males were crossed to five wild-type females to generate drive heterozygotes. These heterozygotes were then crossed to either wild-type or each other (intercross). After bloodfeeding, females were moved into single oviposition cups each containing a smaller plastic cup inside, with water and filter paper for inducing egg laying. The number of eggs and hatched larvae were counted to assess the egg viability.

### Genotyping

Genomic DNA from mosquitoes was extracted using DNAzol Reagent (Invitrogen). Genomic integration of the TARE construct was confirmed by PCR using Q5 High-Fidelity DNA Polymerase (New England Biolabs), and the products were further verified by Sanger sequencing. The targeted *hairy* region was amplified using primers 565 (5’- tcgtatcaacaactgtctgaacgagctg-3’) and 39 (5’-tgaaggatggccggccaatc-3’), yielding a 557 bp product covering all gRNA target sites.

### Cage experiment

Six cage populations (Cage A-F) were established with different initial release ratios. For Cage A-D, homozygous TARE mosquitoes were crossed with wild-type individuals to generate heterozygous eggs, which were pooled with wild-type eggs at ratios approximating intended release levels (thus representing a heterozygous release). For Cage E and F, TARE homozygous eggs were mixed directly with wild-type eggs (representing a homozygous release). The second or third instar larvae were phenotyped to confirm the release ratios, indicated by the percentage of larvae carrying green fluorescence (TARE allele). Larvae were then distributed into trays (approximately 400 larvae each) to standardize density. Pupae were collected daily and transferred to cages. Nearly all adults emerged within one week into a common cage after the first one eclosed, and they were all blood-fed ten days after the first one eclosed. Eggs were collected for 24 h on day 3 post-bloodmeal, and ∼1000 larvae per generation were randomly selected for phenotyping and propagation.

### Maximum-likelihood based fitness analysis of cage populations

Drive carrier frequencies collected across multiple generations and replicate cages were analyzed using a previously published maximum-likelihood inference framework to interpret cage performance, with a simplifying assumption of a single gRNA at the drive target site. The model was implemented in R (v4.0.3) and is available on GitHub (https://github.com/jchamper/Anopheles-TARE-Drive).

In the model, germline cutting of the target site converts the wild-type allele into either a functional resistance allele (R1) or a non-functional resistance allele (R2). No homing-based drive conversion was assumed; instead, drive propagation relies on disruption of wild-type and subsequent elimination of R2 alleles.

Three combinations of germline and embryonic cut rates were evaluated, with or without the formation of R1 alleles. For each parameter set, three alternative sources of fitness cost were independently modeled: viability, combined male mating fitness/female fecundity (with the same parameter for each), and only female fecundity. Models were assessed using the corrected Akaike information criterion (AICc). Models with lower AICc values are considered more likely to explain the observed cage dynamics without overfitting. Parameter estimates from the best-supported models were subsequently used in further modeling analyses.

### Experimental fitness assessment

To further evaluate the fitness of the TARE homozygotes, we recorded the egg number, hatch rate, and life span of this line, with comparison to wild-type control. Eggs from both strains were pooled and reared to the pupal stage. Pupae were screened by fluorescence and set up in separate cages, where homozygotes or wild-types were intercrossed. This procedure was repeated for two generations to minimize batch effects. In the test generation, individual females were placed in paper cups after a blood meal for oviposition. The numbers of eggs and hatched larvae were subsequently recorded for calculating the hatch rate. Newly hatched larvae were collected on the same day and reared in small pots for life span assessment. Three biological replicates were set up per strain, each containing 30 larvae. Developmental stage and survival were monitored daily, and pupae were transferred to paper cups to allow emergence of adults.

### Statistical analysis

To compare the drive inheritance rate, pupation rate, and eclosion rate among groups, a previously reported batch effect analysis method (Champer et al. 2020e, Du et al. 2024, Chen et al. 2024, Faber et al. 2024, Xu et al. 2025, 2026) was used. This model was built based on R program, incorporating variability between cross batches. Briefly, a generalized linear mixed model fitted by maximum likelihood (Adaptive Gauss–Hermite Quadrature, nAGQ = 25). z-tests in this model were applied to calculate *p* values. To compare the egg numbers produced by single female, t-test or one-way ANOVA was applied using Graphpad Prism.

### Modeling

A previously developed SLiM(Haller and Messer 2023) model, which simulated various life stages of mosquitoes(Champer et al. 2022, Liu et al. 2023, Zhu et al. 2024), was utilized to analyze the population dynamics of our TARE drive. Simulations were run for 100 weeks post release, with 3.167 weeks representing one generation in this model. The wild-type population was allowed to equilibrate for 10 generations before releasing drive carriers.

A 4-week homozygote release scheme was modeled with different release ratios and the presence or absence of R1 functional resistance allele formation. Outputs included weekly drive allele frequency, drive carrier frequency, R1 functional resistance allele frequency, and R2 disrupted allele frequency in each week. Other parameters were set based on our experimental data.

Each simulation was independently run five times, and data was collected for figure generation with Python. The default parameters for both drives are listed in Table S1, and corresponding SLiM scripts and data can be found in Github (https://github.com/jchamper/Anopheles-TARE-Drive).

## Results

### Construction of a TARE drive in *Anopheles stephensi*

We aimed to construct a Toxin-Antidote Recessive Embryo (TARE) modification drive in *A. stephensi* by targeting the haplosufficient gene *hairy* (Figure 1A). A proof-of-concept for this drive has been established in *D. melanogaster*, where the drive exhibited high transmission through female heterozygotes and achieved spread in cage populations(Champer et al. 2020c, Metzloff et al. 2022). In this design, the drive incorporates a recoded rescue element for the target gene. Thus, only individuals with at least one drive allele or intact wild-type allele are viable, whereas offspring carrying two disrupted alleles are not (Figure 1B). Successful population modification therefore relies primarily on achieving efficient germline cleavage, if the toxin-antidote mechanism functions as intended. Repair can be by homology-directed repair or end-joining, as long as the sequence becomes disrupted. Although maternal deposition of Cas9 can further enhance inheritance in females, it is not strictly required for drive propagation, with germline cleavage from both males and females always contributing to overall drive increase at the population level(Champer et al. 2020b).

**Figure 1.**
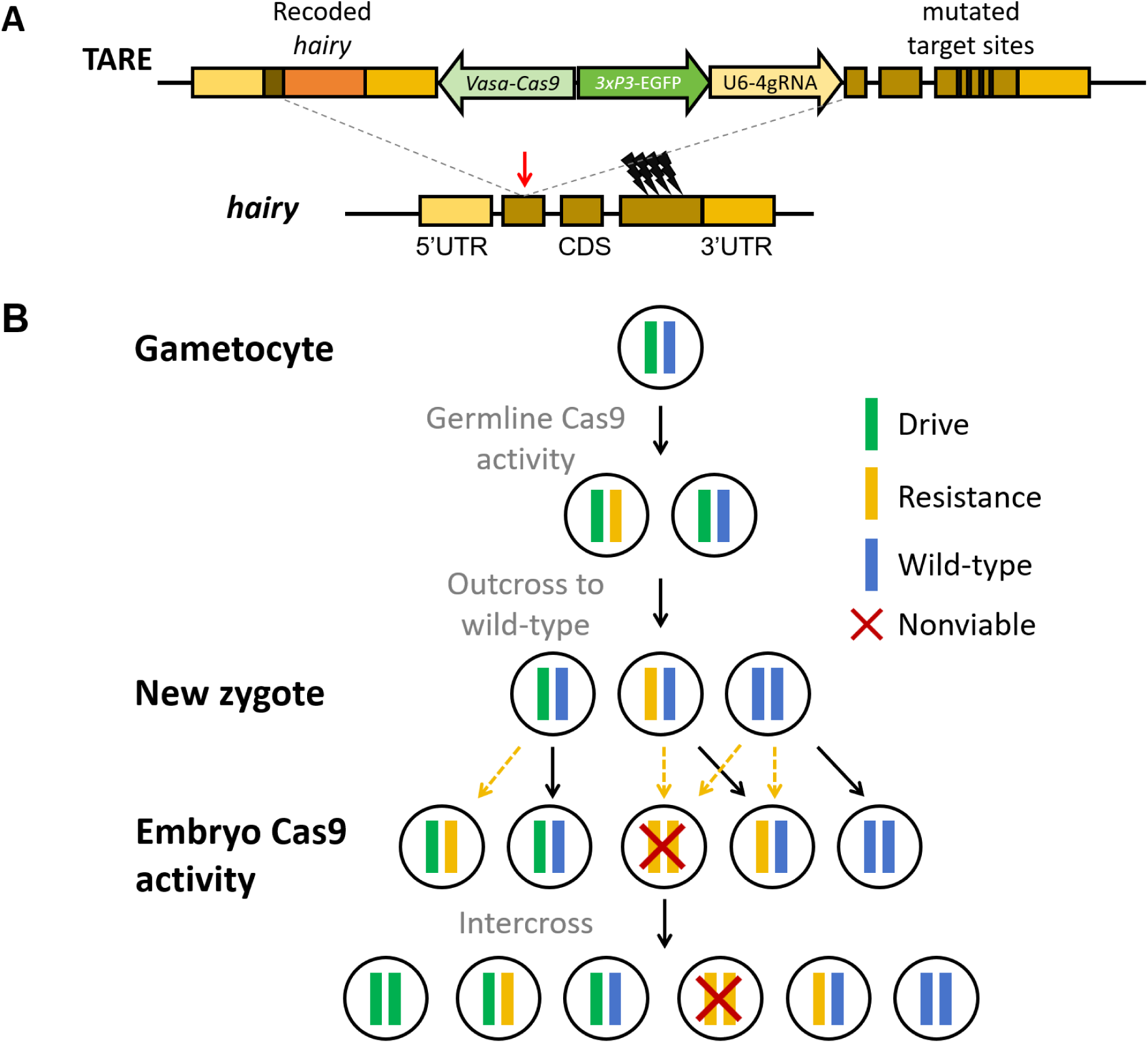
Schematic illustration of the TARE modification gene drive. (**A**) The TARE construct is inserted into the *hairy* coding region. The drive harbors a recoded version of *hairy* terminated with its native 3′ UTR, a 3xP3-EGFP-SV40 marker, and a ribozyme-gRNA cassette consisting of four gRNAs targeting *hairy* for gene disruption. Black lightening symbols indicate gRNA target sites, and a red arrow shows the insertion site. The TARE donor construct contains a modified right homology arm where gRNA target sites were mutated to avoid cleavage of the drive plasmid. (**B**) Cas9/gRNA expressed from the drive allele is expected to cut the wild-type allele in the germline, followed by homology-directed repair or end-joining mediated disruption of target gene. In the offspring of the drive (assumed to mate with wild-type), maternal deposition of Cas9/gRNA is also expected to cleave and disrupt wild-type alleles in the embryo. Offspring inheriting two disrupted alleles are non-viable.

In adapting the toxin-antidote strategy to *A. stephensi* for population modification, we implemented a complete drive design, housing both Cas9 and gRNA expressing cassettes. Notably, the gRNA target sites are ∼780 bp away from the insertion site. Minimal mutations were introduced into the right homology arm at the gRNA target sites to avoid cutting of donor construct, but initial sequencing result revealed that these mutations were not present after integration of the drive into the mosquito genome by homology-directed repair. Morphological analysis of the TARE line did not reveal any observable differences compared to wild-type mosquitoes (except for EGFP fluorescence).

### Drive efficiency in the *Anopheles* TARE line

After obtaining the TARE line, a drive efficiency test was conducted by crossing TARE drive line males (the progeny of injected individuals) with wild-type. Drive heterozygotes were then crossed to wild-type, and offspring were phenotyped. The first batch of data, which was previously reported(Xu et al. 2025), showed that the drive inheritance rate among progeny of drive males (47.3%±1.0%) was significantly lower than the 50% expectation (*p*=0.008, z-test), though the magnitude of this difference was low. It may have been caused by a small viability fitness cost. No significant difference (*p*=0.3237, z test) was observed in drive females compared to the Mendelian expectation for drive inheritance (51.3%±1.3%). In a cross of drive males and drive females, the drive inheritance rate was 71.1%±1.1%, lower than the expected 75% (*p*<0.0001, z-test) (Figure 2A, Data Set S1). Sanger sequencing of non-drive offspring from the intercross of TARE heterozygotes indicated a 12.5% germline cut rate, assuming that males and females have the same cut rate and that maternal deposition was negligible (Table S1).

**Figure 2.**
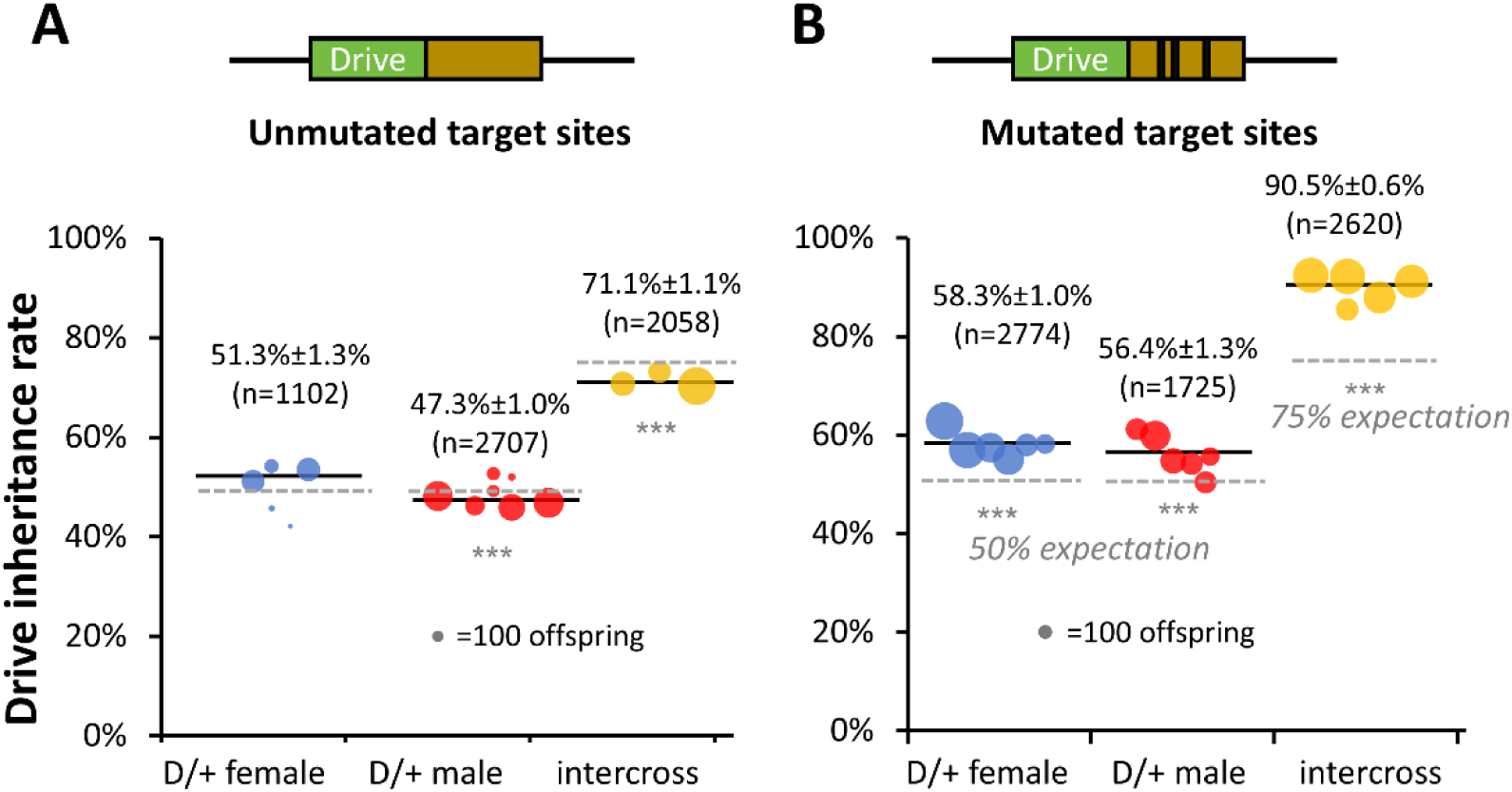
Drive efficiency assessment. Diagrams illustrate the TARE lines tested in two batches where **(A)** target sites in the drive carrier were not mutated [data reported previously (Xu et al. 2025)], and **(B)** with mutated target sites adjacent to the drive allele (which was generated in this study). Drive heterozygotes (D/+) were either outcrossed to wild-type or intercrossed, and their offspring were screened for EGFP fluorescence as a measure of drive inheritance. The percentage of larval progeny with the drive and the total number of larvae screened (n) are shown. Dot size represents the number of progeny from each batch. Mean inheritance rates ± standard error of the mean are shown. Mendelian expectations are shown with light grey dashed lines.Significant differences between drive inheritance rate and Mendelian expectation are marked with *** (*p*<0.001, z-test).

However, we hypothesized that the real cut rate might have been higher. We found that most TARE drive alleles had unmodified gRNA target sites. This means that regardless of whether the drive-adjacent target sites or wild-type chromosome allele was cut, the other could act as a template for homology-directed repair, thus restoring the sequence to wild-type after Cas9 cleavage. We thus generated drive homozygotes, which were maintained for several generations. We expected that eventually, mutations would form in some drive allele target sites, which could then be copied by homology-directed repair to other individuals in the drive line. Subsequent sequencing revealed that all gRNA target sites were mutated except for the gRNA3 site.

This line was then outcrossed to wild-type mosquitoes (using drive males) to reevaluate drive efficiency. In this second batch of crosses, the drive inheritance rates among offspring from drive males and drive females were 56. 4% ± 1.0% and 58.3% ± 1.0%, respectively, both exceeding the 50% Mendelian expectation (*p*< 0.001 for both, z-test) and representing an improvement over the inheritance rates observed in the first batch (Figure 2B, Data Set S1). In offspring derived from intercrosses between drive heterozygotes, the drive inheritance rate increased to 90.5% ± 0.6%, significantly higher than the 75% Mendelian expectation (*p*< 0.0001, z-test). The results of the female drive cross are consistent with low but present maternal deposition, and the results of the intercross are consistent with TARE drives in general, but it was unclear why the drive males achieved significantly higher than 50% drive inheritance. Based on the slightly raised drive inheritance in males, it is possible that *hairy* may be partially haploinsufficient (meaning that disrupted allele carrying non-drive progeny of drive males would have a low rate of nonviability, increasing drive inheritance), but this could be due to random parental health effects (inheritance was lower than 50% for the first batch, see Figure 2A) or reduction in sperm fitness from Cas9 cleavage. It is also possible that a small amount of drive conversion (the drive copying mechanism of homing drives) was taking place.

We further collected non-drive offspring from each cross for Sanger sequencing, which indicated that targets were disrupted at an intermediate rate. Specifically, disrupted alleles were detected in 56.3% (9/16) of offspring from drive males, 18.8% (3/16) from drive females, and 31.0% (9/29) from heterozygote intercrosses, likely resulting from germline or early embryonic cleavage events. No significant difference was found between drive males and females for frequency of resistance alleles (*p*=0.0659, Fisher’s Exact test). Note that some individuals from the female cross and more individuals from the intercross would have been disrupted allele homozygotes and thus nonviable, not appearing in this this sequencing test (so actual cleavage rates were likely somewhat higher).

Notably, one intercross offspring carried a homozygous frameshift mutation (−13 bp) at the final gRNA target site, whereas all other disrupted individuals were heterozygous for disrupted and wild-type alleles (Table S2). This homozygous genotype may reflect either a high amplification bias during PCR or an example of homology-directed repair in the early embryo following cleavage of a wild-type allele, using a disrupted allele formed in the germline of one parent as a repair template. Although this allele could represent a functional resistance allele (albeit one that remained wild-type at most sites), this seems unlikely given the frameshift nature of the mutation, making the explanation of PCR bias or another explanation more likely. Based on these sequencing results, we infer that the gRNA3 target site remained unmodified due to an extremely low cleavage rate, whereas the other gRNAs exhibited higher activity (Table S2).

Our results imply that cut rates of our TARE line was still relatively low compared to previous CRISPR toxin-antidote drives in in *D. melanogaster*(Champer et al. 2020c). While this was not unexpected for cleavage from maternal deposition, which was at a medium level in previous studies(Gantz et al. 2015, Adolfi et al. 2020), germline cleavage rates were still suboptimal, and it remained somewhat unclear if biased inheritance was due to removal of nonviable eggs. To check this and confirm that *hairy* was essential in *A. stephensi*, seven dead eggs were collected from the intercross of TARE heterozygotes for sequencing. All of them were mutated at the target sites (no wild-type alleles were present) and did not carry the drive allele. Combined with the fact that we were able to maintain a homozygous TARE line, it is likely that the *A. stephensi hairy* gene has similar function with its homolog in *D. melanogaster*, being essential for development, and the recoded *hairy* in TARE was able to rescue gene function.

### Reduced egg viability shows TARE drive mechanism

To investigate the mechanism behind the drive bias, we examined the egg viability of crosses with drive heterozygotes. Drive heterozygous males were either crossed with wild-types or intercrossed with drive heterozygous females, and egg number laid by single females and corresponding hatch rates were subsequently measured. The egg number laid per female in the heterozygous intercross group (22.3±8.3) was lower than the heterozygous male x wild-type female group (68.5±6.8) and the wild-type control (51.0±7.1), but no significant difference was detected (p=0.1392 and p=0.2582, respectively, One-way ANOVA). In comparison, hatch rates were markedly reduced when drive heterozygotes were intercrossed (62.3%±9.0%), compared with heterozygous male x wild-type female group (*p*=0.0014, z-test) and wild-type controls (*p*=0.0006, z-test) (Figure 3, Data Set S2). This reduced egg viability supports the notion that our TARE drive operates at least mostly by disrupting wild-type alleles, which are then removed in nonviable individuals.

**Figure 3.**
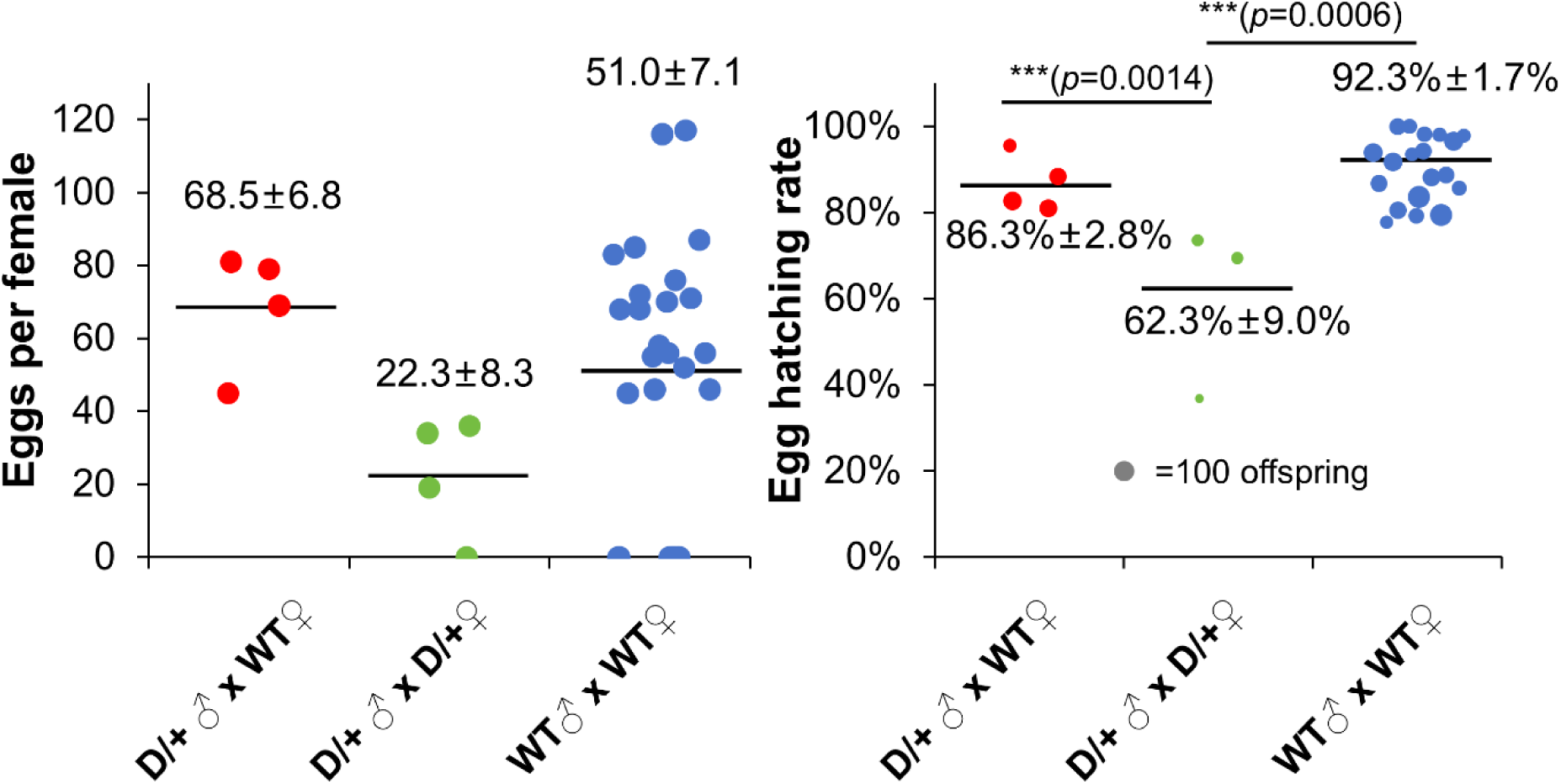
Egg viability assessment of drive heterozygotes. Drive heterozygous males were either crossed to wild-type females or heterozygous females. As a control, wild-type males and females were crossed together. The mean and standard error of the mean of each cross group is shown. The size of dots in the right panel is proportional to the total number of eggs in the single pair crosses. All significant differences are labeled as *** (z-test, *p*<0.001) (eggs per female shows no significant differences, One-way ANOVA).

### Ribozyme-gRNA expression system in *Drosophila*

In the drive efficiency assay described above, the low cleavage rate observed in our *Anopheles* TARE line may reflect insufficient gRNA expression generated by the ribozyme-gRNA system. Proof-of-principle for ribozyme-mediated multiplexing of gRNAs has been demonstrated(Gao and Zhao 2014). However, it remained unclear how this system functions in the context of gene drive. Given the suboptimal performance observed in *Anopheles* mosquito, we considered the possibility that this system might function inefficiently for homing in insects compared to our previously used tRNA system in *D. melanogaster*. To test this hypothesis, we modified a published gRNA construct targeting *yellow-G*(Yang et al. 2022) by replacing the original tRNA-gRNA cassette with a ribozyme-gRNA cassette while retaining the same four gRNAs. The resulting transgenic line exhibited inheritance rates of 82.2% ± 1.4% in drive females and 82.2% ± 2.0% in drive males, with no significant differences compared with the original tRNA-gRNA design (*p*=0.1193 and *p*=0.0705 for females and males, respectively, z-test) (Figure S1, Data Set S3).

These results indicate that the ribozyme-gRNA system is capable of driving robust gRNA expression in the germline of *Drosophila*, achieving performance comparable to the tRNA-gRNA design. However, it is possible that both systems reduce expression, but with the strong *D. melanogaster* U6:3 promoter driving the gRNAs, overall activity remains high. For other species or promoters with lower activity, the use of both ribozymes and tRNAs for gRNA expression may still compromise Cas9 cleavage rates, particularly since the vasa-Cas9 in this TARE line here was able to achieve 100% efficiency when crossed with a line using individual gRNA promoters without ribozyme or tRNA multiplexing(Xu et al. 2025).

### *Anopheles* TARE drive dynamics in cage populations

To evaluate the capacity of our TARE line to spread in a population, we established six cage populations and used different release strategies. Heterozygous TARE individuals were introduced into Cages A through D, and homozygous TARE individuals were released into Cages E and F. The initial release ratios in Cages A and D were 100% and 74%, respectively (Figure 4, Data Set S4). In both cages, the drive carrier frequency increased during the first few generations (after an initial 1-generation drop in cage A), indicating that the TARE drive was functional and able to spread in populations. However, declines were observed beginning at generation 6, suggesting that functional resistance alleles may have appeared with a fitness advantage over the drive. The idea of significant fitness costs for drive is also supported by the weaker performance observed in Cages B, C, E, and F, all of which started with <50% release ratios and declined substantially in just a few generations.

**Figure 4.**
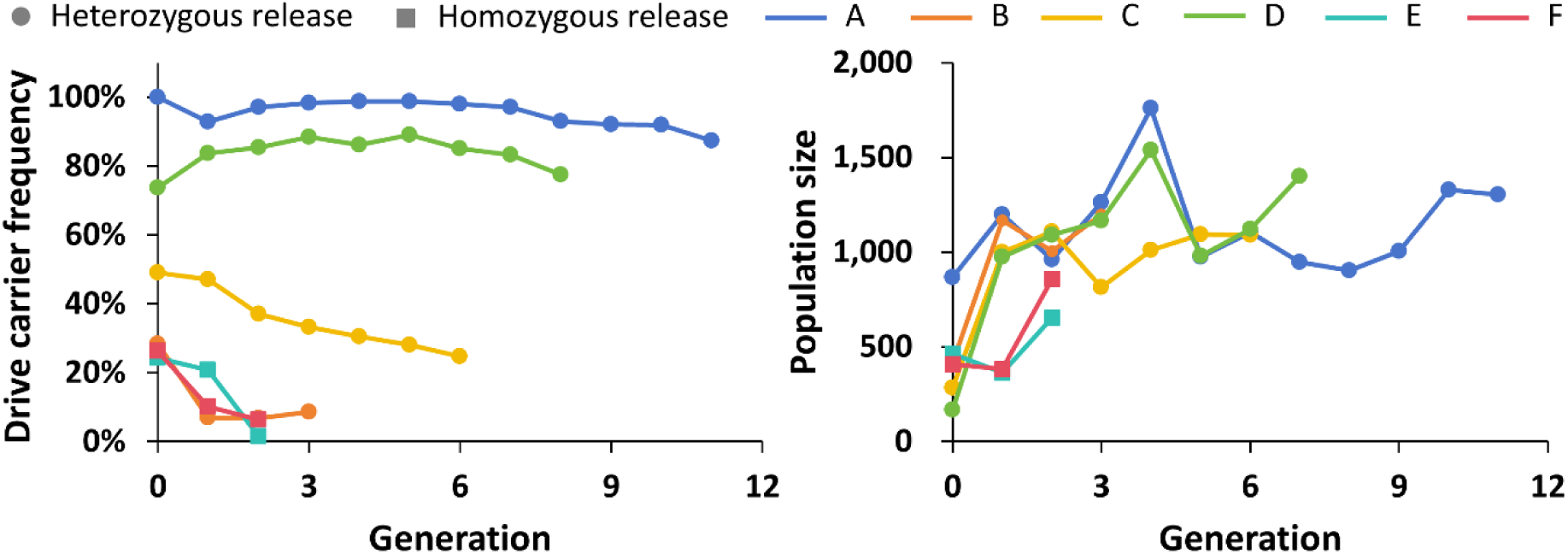
Cage performance of TARE with various release ratios. Six population cages are shown with different colors (marked as A to F). The drive carrier frequency and population size are displayed for each generation. Heterozygous mosquitoes were released into Cage A to Cage D, while homozygous mosquitoes were released into Cage E and Cage F.

Note that a TARE drive without fitness costs does not have an introduction threshold (below which the drive decreases), but one appears only if there are fitness costs. Thus, without fitness costs, all cages would have been expected to increase in frequency until reaching an equilibrium with functional resistance alleles.

To further analyze cage performance, a maximum likelihood analysis was conducted using different combinations of cut rates and fitness parameters (Table S3). These were based on our individual crosses, matching the inheritance rates observed. In one model, germline cleavage with 0.75, while embryo cleavage was 0.35. However, this appeared to have a high rate of embryo cleavage for the intermediate level of germline cleavage, and it could not explain the above-Mendelian inheritance rate in males. Another model had a germline cut rate of 0.78, and embryo cut rate of 0.11, and a fitness multiplier (affecting larval survival) of 0.7 for any individuals with a disrupted allele. A final model dropped this fitness multiplier while keeping the same germline and embryo cut rates. Each of these was evaluated for combinations of fitness type and absolute level of functional resistance allele formation (that could form only in the germline).

Results indicated that cage dynamics were less likely to be driven by viability fitness costs, but costs affecting fecundity and mating choice were likely present (Table S3). Fitness for homozygotes was about 0.5-0.6 (wild-type fitness = 1), and some functional resistance allele formation was present, forming in the germline at a rate of around 1%. However, while the 95% confidence intervals for fitness did not overlap 1, confirming a fitness cost, the 95% confidence interval for functional resistance spanned the entire parameter range. This is likely because the model was overall a poor match for the population cages, indicated by low inferred effective population sizes. This indicates that other unknown factors may have been affecting cage dynamics (such as different fitness in the first generation of released individuals), despite confirmation of a fitness cost. Functional resistance allele formation was also likely to be above zero because otherwise, the high release level cages would have been expected to reach 100% drive carrier frequency.

We hypothesized that with four gRNAs (of which at least three were likely active) and a moderately conserved target site, functional resistance from end-joining was highly unlikely to form in a small laboratory population. However, in a TARE drive, it is possible for the rescue 3′ UTR to be used as a template for homology-directed repair, which might copy only the rescue element(Champer et al. 2020c). Though this did not occur at a detectable level in our previous construct in *D. melanogaster*, *Anopheles* germlines have higher rates of homology-directed repair, and slightly fewer nucleotides of end resection would be needed in this *A. stephensi* version from the cut sites to the native 3′ UTR compared to the *D. melanogaster* version (though still several hundred nucleotides).

To directly assess this possibility of functional resistance alleles in our cages, we randomly genotyped non-fluorescent larvae from the 11th generation of cage A. These larvae were indeed found to all carry a partial drive allele sequence containing only the recoded *hairy* gene and its 3’ UTR, rather than only intact wild-type or fully disrupted alleles (Figure S2). These represent R1 functional resistance. To assess the frequency of R1 allele formation in the germline, we randomly genotyped non-drive larvae derived from crosses of drive heterozygous and wild-type mosquitoes. The R1 frequency was 6.7% (1/15) in both the progeny of drive males crossed to wild-type and drive females crossed to wild-type.

Aside from homology-directed repair, this outcome could also be explained by recombination between the 3′ UTR downstream of the recoded *hairy* rescue and the native *hairy* 3′ UTR adjacent to the drive construct, resulting in loss of Cas9, gRNA, and the marker sequences. However, we consider this less likely due to our homozygous TARE lines remaining stable over several generations, and always producing all drive offspring when crossed with wild-type. Collectively, these results suggest that homologous or repetitive sequences within the drive allele introduced instability and impaired ideal drive function.

### Fitness costs of TARE homozygotes

To assess potential fitness costs associated with our TARE strain, we first conducted a series of experiments examining female fertility and lifespan across larval, pupal, and adult stages. The number of eggs, egg hatch rate, pupation rate, and pupal eclosion rates showed no significant difference between TARE homozygotes and wild-type (Figure S3, Data Set S5). To evaluate lifespan, newly hatched larvae were randomly collected, including 30 individuals in each of three biological replicates, and data from all three replicates were combined for analysis. The larval stage lasted 7-9 days in the TARE strain and 8-10 days in wild-type mosquitoes, while the pupal stage spanned 3-5 days in both groups. Adult longevity was approximately 29-32 days in TARE mosquitoes and 30-31 days in wild-type controls. No apparent differences in survival were observed between the two strains until approximately day 20 (corresponding to ∼10 days post-eclosion), after which TARE adults exhibited a more rapid decline in survival than wild-type mosquitoes (Figure S4, Data Set S6). Notably, in our cage experiments, pupae were collected over a one-week period and adults were allowed to mate for three more days before blood-feeding. Therefore, the majority of mosquitoes remained alive throughout the cage trials and likely all or nearly all mated. This makes it less likely for the reduction in adult longevity to influence the drive inheritance in cage populations, except for the reduced larval time, which would have been an advantage to the TARE drive. This was probably a batch effect, though, and may not have represented a real difference between the lines. The reduced longevity of the TARE line may be indicative of underlying health effects that could well have influenced the cage populations.

### Modeling of *Anopheles* TARE drive dynamics

Although our cage experiments showed issues with the drive, if functional resistance can be overcome, it may still be effective, despite the fitness cost. We thus performed simulations of TARE releases using a mosquito-specific population model and drive performance parameters based on our experimental results (Table S4). Because TARE is a frequency-dependent drive and because a large fraction of the population is at juvenile stages at any moment, we modeled weekly adult releases over a four-week period and tracked the drive allele and carrier frequency.

When R1 functional resistance alleles emerged, the drive frequency declined after initial releases pushed it to a high level (Figure 5 and S5). This was initially driven by replacement of released drive adults with juveniles, where drive frequency was lower. After about 50 weeks, though, functional resistance alleles reached a high rate, and the drive frequency declined more rapidly, reducing the fraction of the mosquito population that was protected from malaria transmission. Assuming functional resistance can be overcome, we found that at the lower release ratio, drive frequencies continuously declined. Though the drive frequency was pushed above the introduction threshold by released adults, newly emerging larvae had a lower frequency. When the release ratio was increased, frequencies initially declined but subsequently rebounded, with the drive carrier frequency remaining high and eventually reaching approximately 100% within 100 weeks. While the drive carrier frequency reached approximately 100%, the drive allele frequency increased more slowly toward a lower equilibrium value, as expected from TARE drives with fitness costs(Champer et al. 2020c, 2020b).

**Figure 5.**
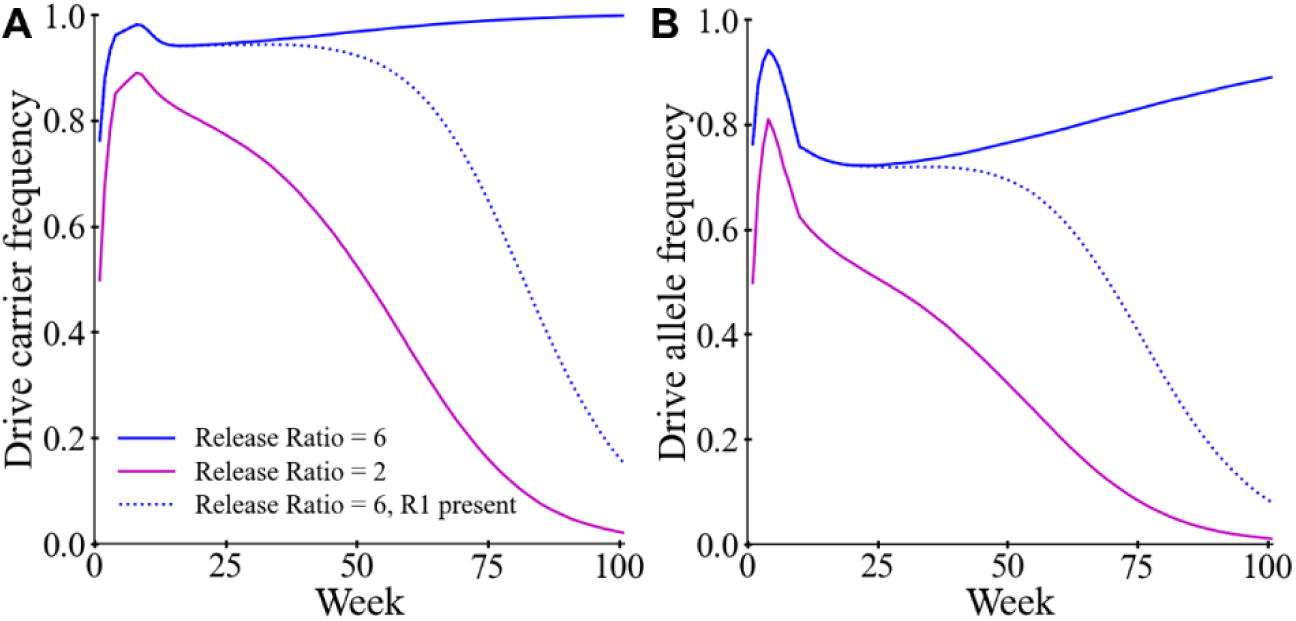
TARE drive performance in the mosquito model. Drive homozygous individuals (both sexes in equal numbers) were released into the population for the first four weeks at the specified release ratio (relative to the size of only the adult female population), with resistance present on one simulation set. Adult (**A**) drive carrier frequency and (**B**) drive allele frequency were monitored each week. See Figure S5 for all allele frequencies for each type of simulation.

## Discussion

Modification gene drives represent a promising strategy for controlling mosquito-borne diseases by spreading anti-pathogen traits while preserving mosquito populations and their ecological roles. Previous studies have demonstrated the feasibility of blocking pathogen transmission either by expressing exogenous antimicrobial effectors or by disrupting endogenous mosquito genes required for pathogen development(Dong et al. 2018, Carballar-Lejarazú et al. 2023, Green et al. 2023, Verkuijl et al. 2025, Habtewold et al. 2025). When coupled to gene drive systems, such traits can spread rapidly through populations and contribute to population level disease reduction(Carballar-Lejarazú et al. 2023, Li et al. 2025). However, most existing modification drives rely on homing-based inheritance bias, which raises concerns about uncontrolled spread beyond target populations and potential failure caused by resistance formation. Consequently, more confined modification drives, which are in theory more refractory to end-joining induced resistance(Champer et al. 2020b), are desirable for future field deployment. The Toxin-Antidote Recessive Embryo (TARE) drive is perhaps the type of confined drive that can be constructed most readily.

In this study, we developed a TARE modification drive in the major malaria vector *Anopheles stephensi* targeting the endogenous gene *hairy*. In *D. melanogaster*, *hairy* is a haplosufficient yet essential gene required for early embryonic segmentation and patterning, serving as a good TARE target(Champer et al. 2020c). Although the function of *hairy* has not been characterized in *Anopheles* mosquitoes, its conserved role across insects suggests a similar role and properties. Consistent with this expectation, our genotyping of nonviable embryos indicates that *hairy* null alleles are also recessive lethal in *A*. *stephensi*. Furthermore, the successful maintenance of a homozygous TARE line in the laboratory demonstrates that the recoded *hairy* sequence within the drive allele is capable of rescuing the endogenous gene. Egg viability was significantly reduced in crosses between drive heterozygotes compared with those involving a drive male only or wild-type controls, as expected in the TARE drive mechanism.

A major issue with our drive was the formation of functional resistance by the recoded rescue element of the drive. This was seen at low rates in a homing drive *in D. melanogaster*(Faber et al. 2025), but never before in a TARE drive. In our construct, identical *hairy* 3′ UTR sequences are present downstream of both the rescue element and the native genomic region adjacent to the drive insertion site. The most likely explanation for functional resistance allele formation is that end resection proceeded enough from the drive target site to reveal the native 3′ UTR, which then matched with the drive 3′ UTR in the rescue element, copying only the rescue element to the cut chromosome. A similar mechanism could also have caused a small amount of drive conversion (copying of the drive) as in homing drives. Another possible explanation is structural instability of the drive allele caused by internal sequence repetition. Such sequence identity could facilitate recombination, leading to partial drive conversion or loss of functional elements. Because this did not affect the stability of long-term TARE lines, it would likely still take place during the process of homology-directed repair, just on the drive allele itself rather than the cleaved wild-type allele. It is known that such instability can affect drive performance. In a *D. melanogaster* homing drive test, replacing native rescue 3′ UTR sequences with orthologs from closely related species substantially improved drive performance(Hou et al. 2024). Recombination driven by repetitive gRNA arrays has likewise been reported to compromise the spread capacity of other gene drive systems(Green et al. 2023). Overall, it is likely that this issue can be alleviated or eliminated by using a 3′ UTR from a different species(Adolfi et al. 2020, Hou et al. 2024), or even a completely different 3′ UTR (though this could potentially affect rescue, causing fitness costs).

Accurate temporal and spatial expression of the rescue element is critical for the success of toxin-antidote gene drives. In our design, the recoded *hairy* sequence was inserted directly into the native *hairy* locus, allowing its expression to be regulated by endogenous genomic elements. This configuration likely provides more faithful rescue than designs relying on ectopic expression from a distant locus driven by selected regulatory elements(Oberhofer et al. 2019). However, this system showed a heavy fitness cost, and even a previous one in *D. melanogaster* targeting *hairy* showed moderate fitness costs. Fitness loss arising from multiple sources represent a major limiting factor for the propagation of our TARE drive. In individual assays, drive homogyzotes and wild-type mosquitoes showed similar performance, except for longevity, which may be indicative of negative health effects in the drive. This effect may result from incomplete functional rescue by the recoded *hairy* gene carried in the drive, which lacks intronic sequences and contains only recoded coding regions. The removed introns may harbor regulatory elements required for native hairy expression and full gene function. Such fitness costs are expected to be more pronounced in individuals carrying two copies of the drive allele, or especially one drive allele together with a disrupted nonfunctional allele. This hypothesis is supported by maximum-likelihood inference based on cage performance data, in which the best-fitting model identified fitness costs associated primarily with mating success and fecundity. Additionally, a model with reduced fitness in heterozygous carrying a disrupted allele suggests that *hairy* may be partially haploinsufficient in mosquitoes. Therefore, careful consideration of gene structure during recoding of *hairy* will be necessary to improve functional rescue efficiency in future designs. It is also possible that our choice of the *vasa* promoter to obtain higher maternal deposition may have negatively affected the drive via somatic expression, creating a disrupted allele that caused fitness costs in drive heterozygotes (this would only cause fitness costs if the rescue element was not haplosufficient). All these issues could potentially be addressed with a new target gene, and perhaps at least partially by switching to a different Cas9 promoter.

Unlike homing-based gene drives, where end-joining mediated resistant alleles can rapidly halt drive spread by blocking further cleavage, TARE drives do not rely on homology-directed copying and are therefore less vulnerable to resistance formation (Champer et al. 2020c) (though we detected functional resistance alleles, they could have formed similarly or potentially at even higher rates in a similar homing-based rescue drive). Instead, TARE performance primarily depends on achieving sufficiently high germline cleavage rates, with embryonic Cas9 activity further enhancing inheritance bias. Notably, high drive efficiency was observed in homing drive strain targeting *doublesex* (which by itself had moderate activity) when combined with our TARE drive, but not when TARE was deployed alone, indicating that the *vasa*-Cas9 component itself was highly active(Xu et al. 2025). This suggests that the reduced efficiency observed in our standalone TARE line likely arises from other design-specific factors. One plausible explanation is insufficient gRNA expression, leading to suboptimal cleavage in the germline and early embryo. The *A. stephensi* U6 promoter and associated terminal sequences used in this study have been validated previously (Gantz et al. 2015, Adolfi et al. 2020, Xu et al. 2025), suggesting that promoter choice is unlikely to be the limiting factor. Instead, the reduced activity may be caused by low efficiency of the ribozyme-gRNA expression system. To evaluate this possibility, we tested the ribozyme-gRNA design in a homing gene-drive system targeting *yellow-G* in *Drosophila* and observed gRNA activity comparable to that of the tRNA-gRNA multiplexing system. It thus remains unclear why the ribozyme-gRNA performance was reduced in *A. stephensi*. It is possible that both tRNA and ribozyme systems reduce activity, meaning that for most species, individual gRNA promoters should be used for each gRNA (unless one is found with extraordinarily high activity, like the U6:3 promoter in *D. melanogaster*). Another possible explanation would be position effects around the *hairy* locus that inhibit gRNA transcription, which could be resolved with a different target site.

Note that this study focused on evaluating a prototype TARE system in *A. stephensi* without incorporating anti-disease effector components. Nevertheless, multiple strategies for blocking malaria transmission in mosquitoes have been demonstrated in previous studies. For example, exogenous expression of anti-*Plasmodium* single-chain antibodies, such as m1C3 and m2A10, has been shown to interfere with parasite development in *Anopheles* mosquitoes(Isaacs et al. 2012, Carballar-Lejarazú et al. 2023). In addition, complete disruption or targeted modification of endogenous mosquito genes required for *Plasmodium* infection can greatly reduce malaria transmission(Garver et al. 2009, Dong et al. 2018, Habtewold et al. 2025). These approaches could be incorporated into future improved TARE systems to generate release candidates with intrinsic transmission-blocking capacity.

Overall, our study provides important insights into the design principles required for effective and confined gene drive in the non-model insect *A. stephensi*. The results suggest that high-efficiency TARE drives in this species may be achievable through relatively modest design modifications. These include improving gRNA expression with individual promoters and not tRNAs/ribozymes, reducing sequence identity within the 3′ UTR rescue region, and potentially using a different target gene and/or Cas9 promoter. These refinements could substantially enhance drive stability and performance. Ehen combined with additional of anti-malarial effector genes, they may allow highly efficient but confined population modification gene drives for malaria control.

## Acknowledgements

This study was supported by grants from the National Natural Science Foundation of China (32302455, 32270672, and W2432018), the Beijing Natural Science Foundation (IS24039), and the Center for Life Sciences. Xuejiao Xu was supported in part by the Postdoctoral Fellowship of Peking-Tsinghua Center for Life Sciences.

## Supplemental Information

**Table S1.** Genotypes of non-drive offspring from the intercross of TARE heterozygotes.

| Genotypes | gRNA1 | gRNA2 | gRNA3 | gRNA4 | Percentage |
| --- | --- | --- | --- | --- | --- |
| WT | WT | WT | WT | WT | 75% (12/16) |
| #1 | R2/WT | R2/WT | WT | R2/WT | 18.75% (3/16) |
| #2 | WT | WT | WT | R2/WT | 6.25% (1/16) |

WT means wild-type genotype, and R2/WT means heterozygous for both resistance and wild-type alleles. #1 and #2 indicate different mutation types, and their frequencies obtained by sequencing are listed in the table.

**Table S2.** Genotypes of non-drive offspring from the intercross of TARE heterozygotes.

| Genotypes | gRNA1 | gRNA2 | gRNA3 | gRNA4 | Number/Total number |
| --- | --- | --- | --- | --- | --- |
| WT | WT | WT | WT | WT | 20/29 |
| #1 | WT | WT | WT | R2(-13 bp) | 1/29 |
| #2 | R2/WT | R2/WT | WT | R2/WT | 3/29 |
| #3 | WT | R2/WT | WT | WT | 1/29 |
| #4 | R2/R2 | WT | WT | WT | 1/29 |
| #5 | R2/WT | WT | WT | R2/WT | 1/29 |

**Table S3.** Default parameters for the mosquito model.

| Item | Default value |
| --- | --- |
| Number of adult females | 50000 |
| Low-density growth rate | 6 |
| Germline resistance allele formation rate | 0.78 |
| Embryo resistance cut rate | 0.11 |
| Drive conversion rate | 0 |
| Drive homozygote fitness | 0.5 |
| Nonfunctional resistance fitness | 0.7 |
| Functional resistance rate | 0.007693414 |

**Table S4.**
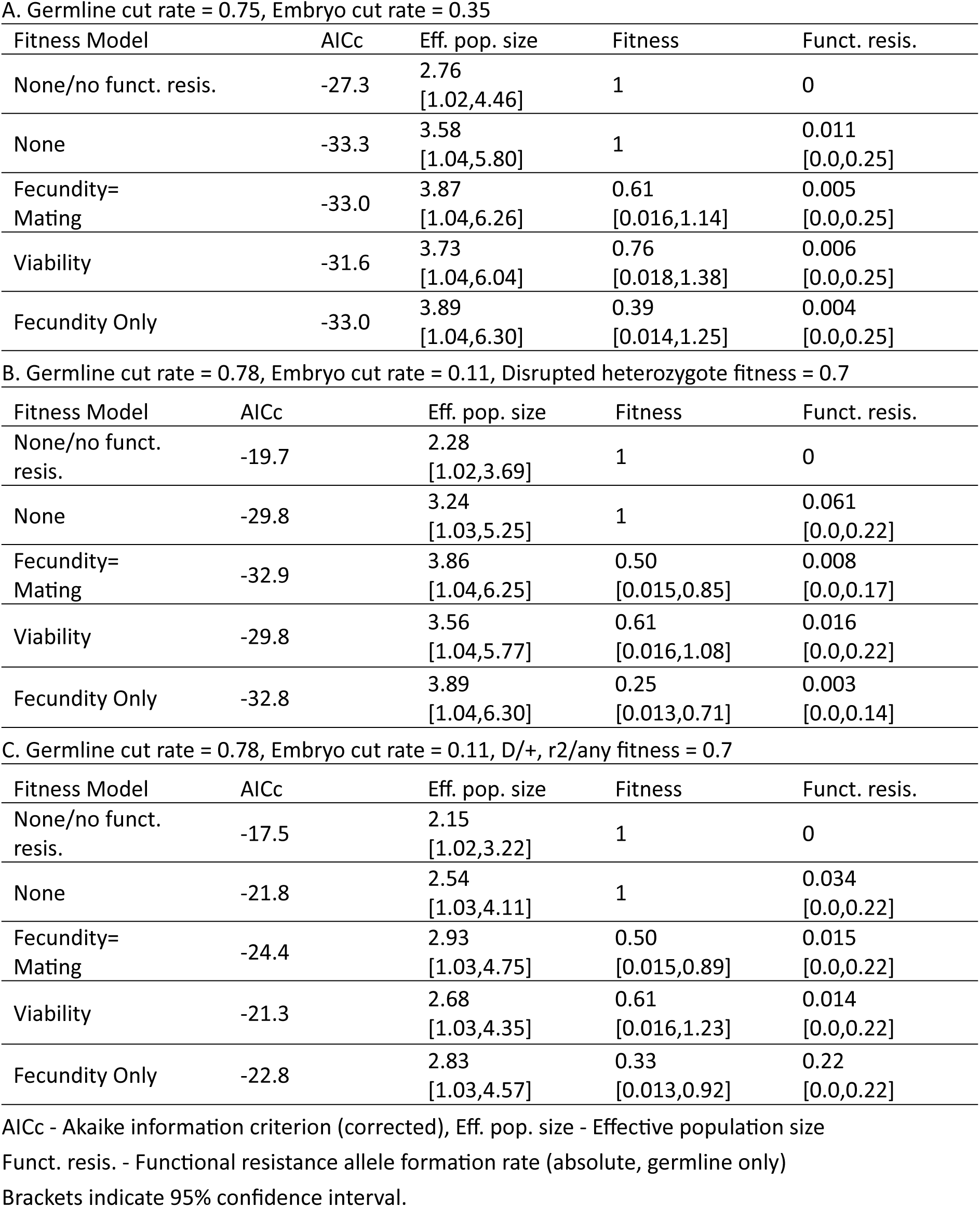
Maximum likelihood analysis of cage experiments.

**Figure S1.**
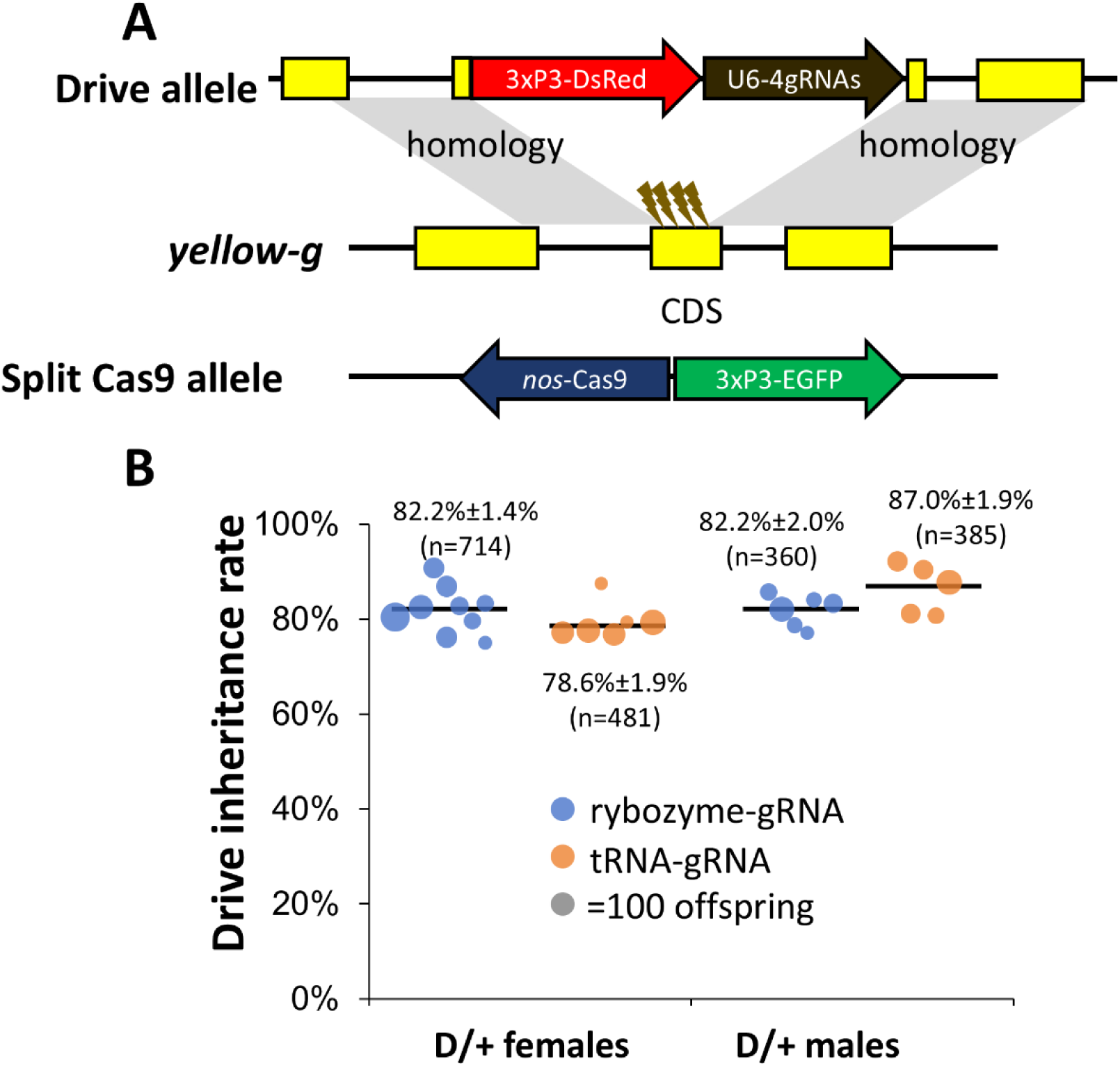
Split gene drive in *D. melanogaster* for testing gRNA expression with ribozyme-gRNA system. (**A**) Schematic of the drive construct. The drive contains a 3xP3-DsRed-SV40 marker and a ribozyme-gRNA cassette that expresses four gRNAs targeting coding sequence of *yellow-g*. A split Cas9 source was provided for drive efficiency testing. (**B**) Drive efficiency test of ribozyme-gRNA construct and a tRNA-gRNA construct reported in the previous study(Yang et al. 2022), identical except for replacement of tRNAs with ribozymes. Drive males were crossed with Cas9 females to generate double heterozygotes, which were crossed to wild-type, and their offspring were phenotyped for drive inheritance. The mean and standard error of the mean (SEM) was displayed. The size of dots represents offspring sample size of a single pair cross. No significant difference was found between tRNA-gRNA and ribozyme-gRNA designs.

**Figure S2.**
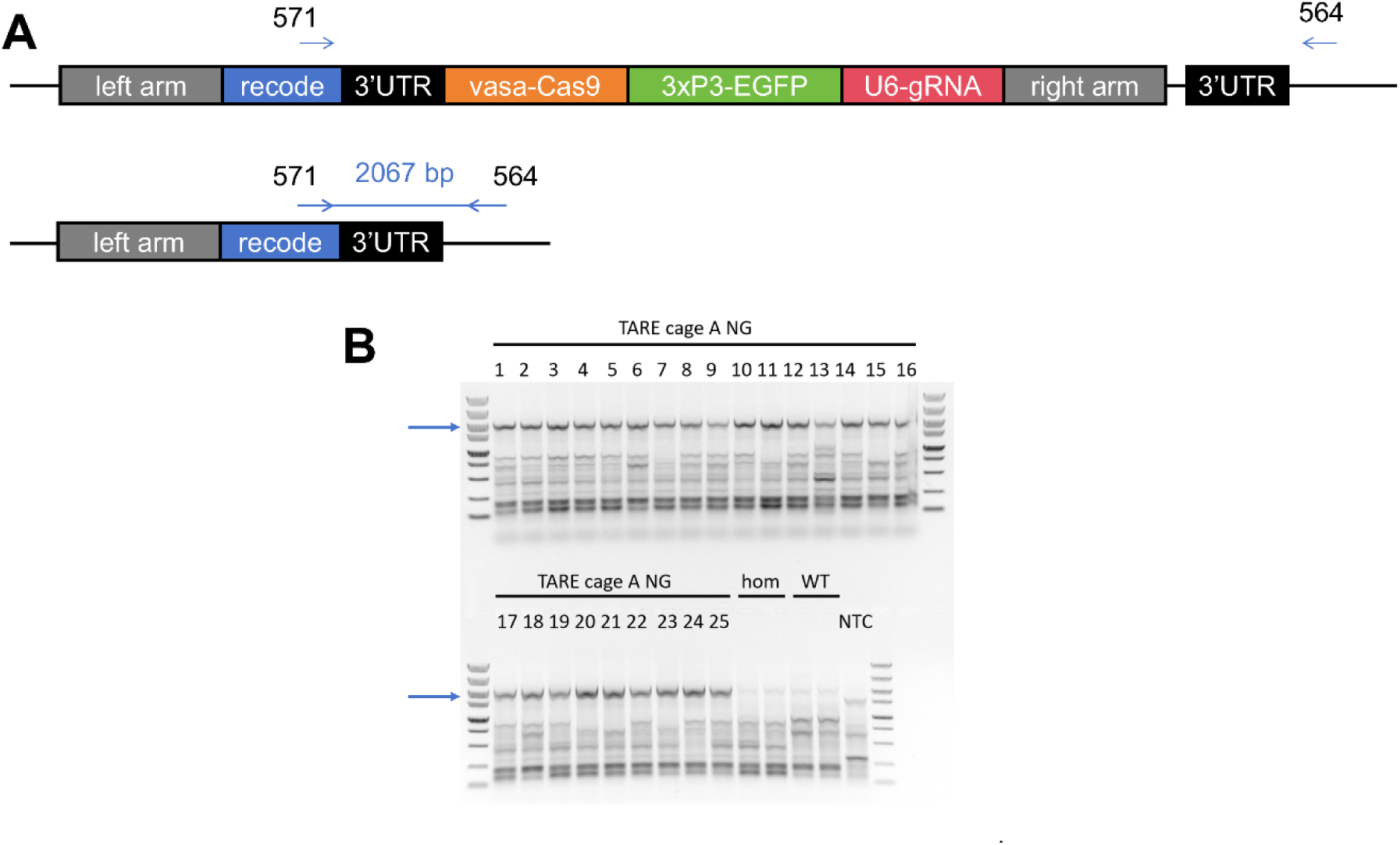
Genotyping of non-fluorescent larvae from Cage A in the 11^th^ generation. (**A**) The upper construct shows a normal drive allele, while the lower one illustrates the non-fluorescent transgene that was the results of undesired homology-directed repair found in Cage A. The locations of primers (571 and 564) used for genotyping were marked. (**B**) In total, 25 larvae were genotyped. The two TARE homozygotes (hom) from the stock cage and two wild-type larvae (WT) were used as negative controls. NTC: no template control. The expected size of undesired homology-directed repair product is marked with blue arrow in the gel image.

**Figure S3.**
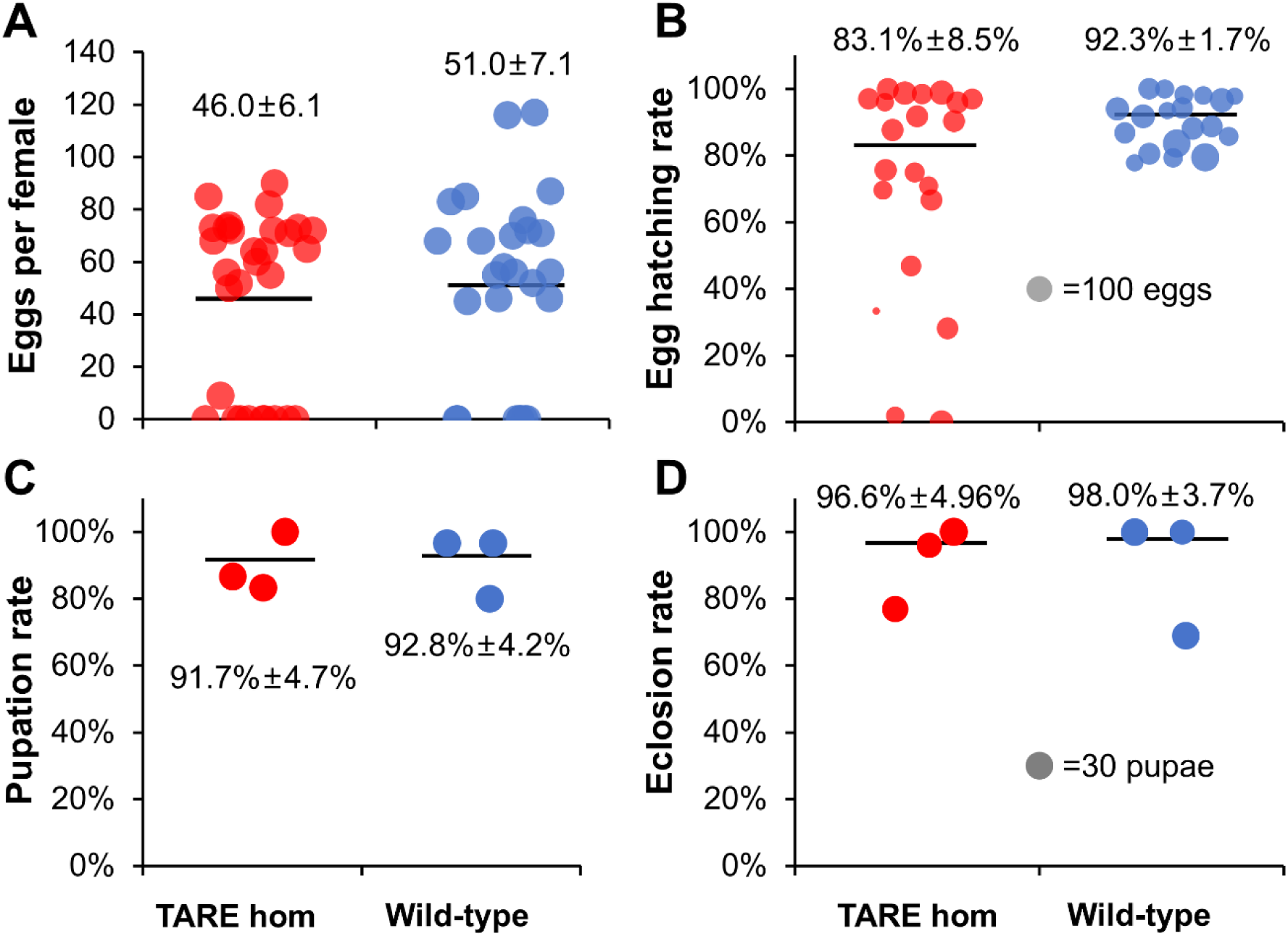
Fitness comparison between TARE homozygotes and wild-type mosquitoes. Four aspects, including (**A**) egg number laid by per female, (**B**) egg-to-larva hatch rate, (**C**) larva-to-pupa pupation rate, and (**D**) pupa-to-adult eclosion rate, were recoded. Mean and SEM were shown in the figure. No significant difference was found for all four parameters by z-test (Egg hatching rate, Pupation rate and Eclosion rate) or t-test (Eggs per female).

**Figure S4.**
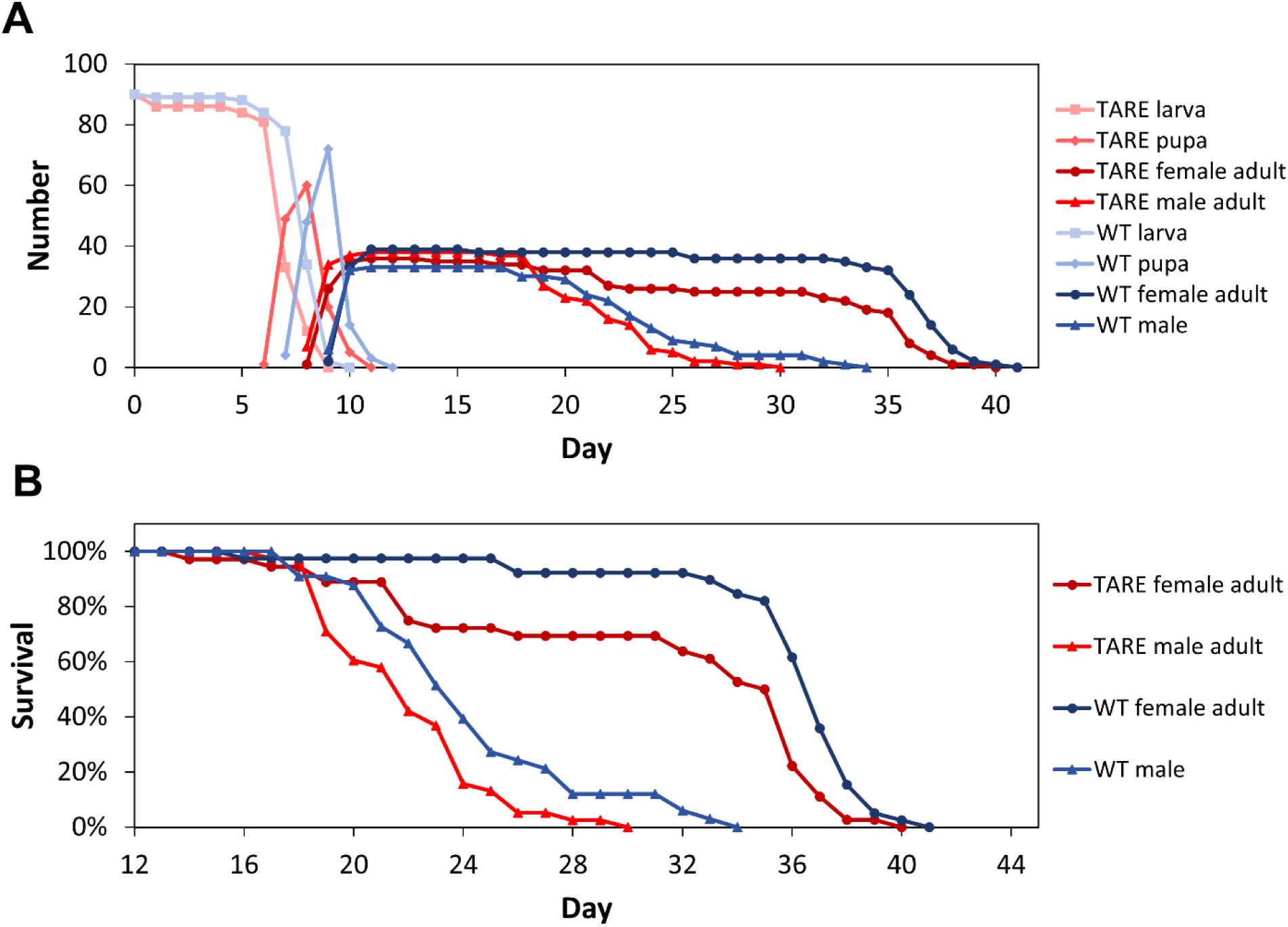
Lifespan comparison between TARE homozygotes and wild-type. (**A**) The total number of mosquitoes at different developmental stages was counted daily, starting with three separate batches of 30 each. Newly hatched larvae were collected on the same day, designated as day 0. (**B**) Female and male adult survival rates were recorded starting from day 12, when all mosquitoes had emerged as adults. The survival rate between two groups was significantly different (*p*=0.0025, Log-rank Mantel-Cox test). Developmental stages are indicated by different colors and symbols.

**Figure S5.**
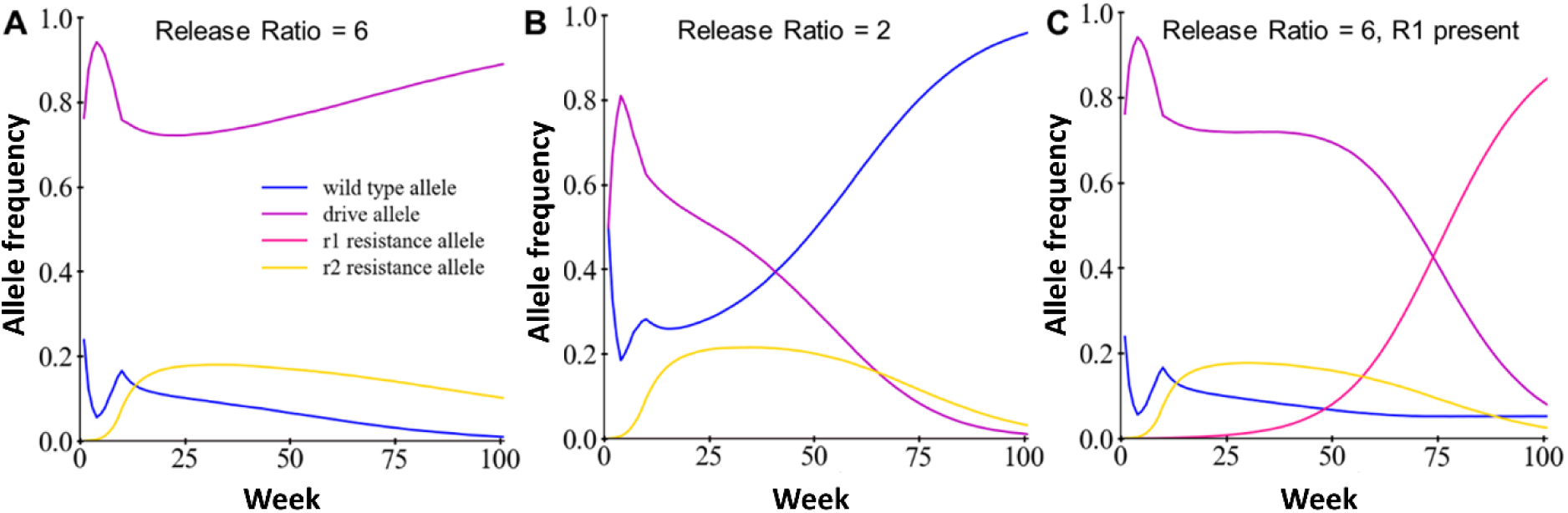
Allele frequencies in TARE drive simulations. Allele frequency data for simulations in Figure 5. Simulations include (**A**) high release ratio, (**B**) low release ratio, and (**C**) high release ratio but with functional resistance allele (R1) formation.

## References

Adolfi, A., V. M. Gantz, N. Jasinskiene, H.-F. Lee, K. Hwang, G. Terradas, E. A. Bulger, A. Ramaiah, J. B. Bennett, J. J. Emerson, J. M. Marshall, E. Bier, and A. A. James. 2020. Efficient population modification gene-drive rescue system in the malaria mosquito *Anopheles stephensi*. Nature Communications 11:5553.

Anderson, M. A. E., E. Gonzalez, J. X. D. Ang, L. Shackleford, K. Nevard, S. A. N. Verkuijl, M. P. Edgington, T. Harvey-Samuel, and L. Alphey. 2023. Closing the gap to effective gene drive in *Aedes aegypti* by exploiting germline regulatory elements. Nature Communications 14:338.

Ang, J. X., S. A. Verkuijl, M. A. Anderson, and L. Alphey. 2025. Synthetic homing endonuclease gene drives to revolutionise *Aedes aegypti* biocontrol — game changer or pipe dream? Current Opinion in Insect Science 70:101373.

Bier, E. 2022. Gene drives gaining speed. Nature Reviews. Genetics 23:5–22.

Carballar-Lejarazú, R., Y. Dong, T. B. Pham, T. Tushar, R. M. Corder, A. Mondal, H. M. Sánchez C., H.-F. Lee, J. M. Marshall, G. Dimopoulos, and A. A. James. 2023. Dual effector population modification gene-drive strains of the African malaria mosquitoes, *Anopheles gambiae* and *Anopheles coluzzii*. Proceedings of the National Academy of Sciences of the United States of America 120:e2221118120.

Champer, J., S. E. Champer, I. K. Kim, A. G. Clark, and P. W. Messer. 2020a. Design and analysis of CRISPR-based underdominance toxin-antidote gene drives. Evolutionary Applications:eva.13180.

Champer, J., I. K. Kim, S. E. Champer, A. G. Clark, and P. W. Messer. 2020b. Performance analysis of novel toxin-antidote CRISPR gene drive systems. BMC Biology 18:27.

Champer, J., E. Lee, E. Yang, C. Liu, A. G. Clark, and P. W. Messer. 2020c. A toxin-antidote CRISPR gene drive system for regional population modification. Nature Communications 11:1082.

Champer, J., E. Yang, E. Lee, J. Liu, A. G. Clark, and P. W. Messer. 2020d. A CRISPR homing gene drive targeting a haplolethal gene removes resistance alleles and successfully spreads through a cage population. Proceedings of the National Academy of Sciences of the United States of America 117:24377–24383.

Champer, S. E., I. K. Kim, A. G. Clark, P. W. Messer, and J. Champer. 2022. *Anopheles* homing suppression drive candidates exhibit unexpected performance differences in simulations with spatial structure. eLife 11:e79121.

Champer, S. E., S. Y. Oh, C. Liu, Z. Wen, A. G. Clark, P. W. Messer, and J. Champer. 2020e. Computational and experimental performance of CRISPR homing gene drive strategies with multiplexed gRNAs. Science Advances 6:eaaz0525.

Chen, W., J. Guo, Y. Liu, and J. Champer. 2024. Population suppression by release of insects carrying a dominant sterile homing gene drive targeting doublesex in Drosophila. Nature Communications 15:1–13.

Dhole, S., A. L. Lloyd, and F. Gould. 2019. Tethered homing gene drives: A new design for spatially restricted population replacement and suppression. Evolutionary Applications 12:1688–1702.

DiCarlo, J. E., A. Chavez, S. L. Dietz, K. M. Esvelt, and G. M. Church. 2015. Safeguarding CRISPR-Cas9 gene drives in yeast. Nature Biotechnology 33:1250–1255.

Dong, Y., M. L. Simões, E. Marois, and G. Dimopoulos. 2018. CRISPR/Cas9-mediated gene knockout of Anopheles gambiae FREP1 suppresses malaria parasite infection. PLoS Pathogens 14:e1006898.

Du, J., W. Chen, X. Jia, X. Xu, E. Yang, R. Zhou, Y. Zhang, M. Metzloff, P. W. Messer, and J. Champer. 2024. Germline Cas9 promoters with improved performance for homing gene drive. Nature Communications 15:4560.

Faber, N. R., J. Champer, B. A. Pannebakker, B. J. Zwaan, and J. van den Heuvel. 2025. Investigating multiple types of resistance against a homing gene drive in European populations of Drosophila melanogaster. bioRxiv:2025.06.06.658298.

Faber, N. R., X. Xu, J. Chen, S. Hou, J. Du, B. A. Pannebakker, B. J. Zwaan, J. van den Heuvel, and J. Champer. 2024. Improving the suppressive power of homing gene drive by co-targeting a distant-site female fertility gene. Nature Communications 15:9249.

Feng, R., and J. Champer. 2024. Deployment of tethered gene drive for confined suppression in continuous space requires avoiding drive wave interference. Molecular Ecology 33:e17530.

Gantz, V. M., N. Jasinskiene, O. Tatarenkova, A. Fazekas, V. M. Macias, E. Bier, and A. A. James. 2015. Highly efficient Cas9-mediated gene drive for population modification of the malaria vector mosquito *Anopheles stephensi*. Proceedings of the National Academy of Sciences of the United States of America 112:E6736–E6743.

Gao, Y., and Y. Zhao. 2014. Self-processing of ribozyme-flanked RNAs into guide RNAs in vitro and in vivo for CRISPR-mediated genome editing. J Integr Plant Biol 56:343–349.

Garver, L. S., Y. Dong, and G. Dimopoulos. 2009. Caspar Controls Resistance to Plasmodium falciparum in Diverse Anopheline Species. PLOS Pathogens 5:e1000335.

Green, E. I., E. Jaouen, D. Klug, R. Proveti Olmo, A. Gautier, S. Blandin, and E. Marois. 2023. A population modification gene drive targeting both Saglin and Lipophorin impairs Plasmodium transmission in Anopheles mosquitoes. eLife 12:e93142.

Grunwald, H. A., V. M. Gantz, G. Poplawski, X.-R. S. Xu, E. Bier, and K. L. Cooper. 2019. Super-Mendelian inheritance mediated by CRISPR-Cas9 in the female mouse germline. Nature 566:105–109.

Guichard, A., T. Haque, M. Bobik, X.-R. S. Xu, C. Klanseck, R. B. S. Kushwah, M. Berni, B. Kaduskar, V. M. Gantz, and E. Bier. 2019. Efficient allelic-drive in *Drosophila*. Nature Communications 10:1640.

Habtewold, T., D. W. Lwetoijera, A. Hoermann, R. Mashauri, F. Matwewe, R. Mwanga, P. Kweyamba, G. Maganga, B. P. Magani, R. Mtama, M. A. Mahonje, M. M. Tambwe, F. Tarimo, P. R. Chennuri, J. A. Cai, G. Del Corsano, P. Capriotti, P. Sasse, J. Moore, D. Hudson, A. Manjurano, B. Tarimo, D. Vlachou, S. Moore, N. Windbichler, and G. K. Christophides. 2025. Gene-drive-capable mosquitoes suppress patient-derived malaria in Tanzania. Nature:1–7.

Haller, B. C., and P. W. Messer. 2023. SLiM 4: Multispecies Eco-Evolutionary Modeling. The American Naturalist 201:E127–E139.

Hou, S., J. Chen, R. Feng, X. Xu, N. Liang, and J. Champer. 2024. A homing rescue gene drive with multiplexed gRNAs reaches high frequency in cage populations but generates functional resistance. Journal of Genetics and Genomics 51:836–843.

Isaacs, A. T., N. Jasinskiene, M. Tretiakov, I. Thiery, A. Zettor, C. Bourgouin, and A. A. James. 2012. Transgenic *Anopheles stephensi* coexpressing single-chain antibodies resist *Plasmodium falciparum* development. Proceedings of the National Academy of Sciences of the United States of America 109:E1922–1930.

Kyrou, K., A. M. Hammond, R. Galizi, N. Kranjc, A. Burt, A. K. Beaghton, T. Nolan, and A. Crisanti. 2018. A CRISPR-Cas9 gene drive targeting *doublesex* causes complete population suppression in caged *Anopheles gambiae* mosquitoes. Nature Biotechnology 36:1062–1066.

Larrosa-Godall, M., J. X. D. Ang, P. T. Leftwich, E. Gonzalez, L. Shackleford, K. Nevard, R. Noad, M. A. E. Anderson, and L. Alphey. 2025. Challenges in developing a split drive targeting dsx for the genetic control of the invasive malaria vector Anopheles stephensi. Parasites & Vectors 18:46.

Li, Z., Y. Dong, L. You, R. M. Corder, J. Arzobal, A. Yeun, L. Yang, J. M. Marshall, G. Dimopoulos, and E. Bier. 2025. Driving a protective allele of the mosquito FREP1 gene to combat malaria. Nature 645:746–754.

Liu, Y., B. Jiao, J. Champer, and W. Qian. 2024. Overriding Mendelian inheritance in Arabidopsis with a CRISPR toxin-antidote gene drive that impairs pollen germination. Nature Plants 10:910–922.

Liu, Y., W. Teo, H. Yang, and J. Champer. 2023. Adversarial interspecies relationships facilitate population suppression by gene drive in spatially explicit models. Ecology Letters 26:1174–1185.

Metzloff, M., E. Yang, S. Dhole, A. G. Clark, P. W. Messer, and J. Champer. 2022. Experimental demonstration of tethered gene drive systems for confined population modification or suppression. BMC Biology 20:119.

Nolan, T. 2021. Control of malaria-transmitting mosquitoes using gene drives. Philosophical Transactions of the Royal Society B: Biological Sciences 376:20190803.

Oberhofer, G., T. Ivy, and B. A. Hay. 2019. Cleave and Rescue, a novel selfish genetic element and general strategy for gene drive. Proceedings of the National Academy of Sciences of the United States of America 116:6250–6259.

Oberhofer, G., T. Ivy, and B. A. Hay. 2020. Gene drive and resilience through renewal with next generation Cleave and Rescue selfish genetic elements. Proceedings of the National Academy of Sciences of the United States of America 117:9013–9021.

Oberhofer, G., M. L. Johnson, T. Ivy, I. Antoshechkin, and B. A. Hay. 2024. Cleave and Rescue gamete killers create conditions for gene drive in plants. Nature Plants 10:936–953.

Pescod, P., G. Bevivino, A. Anthousi, J. Shepherd, R. Shelton, F. Lombardo, and T. Nolan. 2024. Homing gene drives can transfer rapidly between *Anopheles gambiae* strains with minimal carryover of flanking sequences. Nature Communications 15:6846.

Pfitzner, C., M. A. White, S. G. Piltz, M. Scherer, F. Adikusuma, J. N. Hughes, and P. Q. Thomas. 2020. Progress Toward Zygotic and Germline Gene Drives in Mice. The CRISPR Journal 3:388–397.

Speth, Z. J., D. G. Rehard, P. J. Norton, and A. W. E. Franz. 2025. Performance of two low-threshold population replacement gene drives in cage populations of the yellow fever mosquito, Aedes aegypti. PLoS genetics 21:e1011757.

Verkuijl, S. A. N., J. X. D. Ang, L. Alphey, M. B. Bonsall, and M. A. E. Anderson. 2022. The Challenges in Developing Efficient and Robust Synthetic Homing Endonuclease Gene Drives. Frontiers in Bioengineering and Biotechnology 10:856981.

Verkuijl, S. A. N., G. Del Corsano, P. Capriotti, P.-S. Yen, M. G. Inghilterra, P. Selvaraj, A. Hoermann, A. Martinez-Sanchez, C. V. Ukegbu, T. M. Kebede, D. Vlachou, G. K. Christophides, and N. Windbichler. 2025. A suppression-modification gene drive for malaria control targeting the ultra-conserved RNA gene mir-184. Nature Communications 16:1–14.

Walter, M., A. K. Haick, R. Riley, P. A. Massa, D. E. Strongin, L. M. Klouser, M. A. Loprieno, L. Stensland, T. K. Santo, P. Roychoudhury, M. Aubert, M. P. Taylor, K. R. Jerome, and E. Verdin. 2024. Viral gene drive spread during herpes simplex virus 1 infection in mice. Nature Communications 15:8161.

Wang, G.-H., A. Hoffmann, and J. Champer. 2025. Gene drive and symbiont technologies for control of mosquito-borne diseases. Annual Review of Entomology 70:229–249.

Xu, X., J. Chen, Y. Wang, Y. Liu, Y. Zhang, J. Yang, X. Yang, B. Chen, Z. He, and J. Champer. 2025. Gene drive-based population suppression in the malaria vector *Anopheles stephensi*. Nature Communications 16:1007.

Xu, X., J. Fang, J. Chen, J. Yang, X. Yang, S. Hou, W. Sun, and J. Champer. 2026. Assessing target genes for homing suppression gene drive. The EMBO Journal.

Xu, X., T. Harvey-Samuel, H. A. Siddiqui, J. X. D. Ang, M. E. Anderson, C. M. Reitmayer, E. Lovett, P. T. Leftwich, M. You, and L. Alphey. 2022. Toward a CRISPR-Cas9-Based Gene Drive in the Diamondback Moth *Plutella xylostella*. The CRISPR Journal 5:224–236.

Yadav, A. K., C. Butler, A. Yamamoto, A. A. Patil, A. L. Lloyd, and M. J. Scott. 2023. CRISPR/Cas9-based split homing gene drive targeting *doublesex* for population suppression of the global fruit pest *Drosophila suzukii*. Proceedings of the National Academy of Sciences of the United States of America 120:e2301525120.

Yang, E., M. Metzloff, A. M. Langmüller, X. Xu, A. G. Clark, P. W. Messer, and J. Champer. 2022. A homing suppression gene drive with multiplexed gRNAs maintains high drive conversion efficiency and avoids functional resistance alleles. G3 (Bethesda, Md.) 12:jkac081.

Yang, X., X. Xu, Y. Chen, J. Wei, W. Huang, S. Wu, J. Champer, and J. Wang. 2025. Assessment of drive efficiency and resistance allele formation of a homing gene drive in the mosquito Aedes aegypti. Journal of Pest Science 98:899–911.

Zhu, J., J. Chen, Y. Liu, X. Xu, and J. Champer. 2024. Population suppression with dominant female-lethal alleles is boosted by homing gene drive. BMC Biology 22:201.

